# A junction coverage compatibility score to quantify the reliability of transcript abundance estimates and annotation catalogs

**DOI:** 10.1101/378539

**Authors:** Charlotte Soneson, Michael I Love, Rob Patro, Shobbir Hussain, Dheeraj Malhotra, Mark D. Robinson

## Abstract

Most methods for statistical analysis of RNA-seq data take a matrix of abundance estimates for some type of genomic features as their input, and consequently the quality of any obtained results are directly dependent on the quality of these abundances. Here, we present the junction coverage compatibility (JCC) score, which provides a way to evaluate the reliability of transcript-level abundance estimates as well as the accuracy of transcript annotation catalogs. It works by comparing the observed number of reads spanning each annotated splice junction in a genomic region to the predicted number of junction-spanning reads, inferred from the estimated transcript abundances and the genomic coordinates of the corresponding annotated transcripts. We show that while most genes show good agreement between the observed and predicted junction coverages, there is a small set of genes that do not. Genes with poor agreement are found regardless of the method used to estimate transcript abundances, and the corresponding transcript abundances should be treated with care in any downstream analyses.

## Introduction

High-throughput sequencing of the transcriptome (RNA-seq) is used for a broad range of applications in biology and medicine. Most of these involve comparing expression levels of genetic features (e.g., genes, transcripts or exons) between samples, and the quality of the results from any such study will therefore be directly dependent on the correctness of the expression estimates for the particular features of interest. The ability to obtain accurate estimates, in turn, depends on the quality and quantity of the available data, as well as the completeness and correctness of the utilized reference annotation. In general, reliable abundance estimation is easier to achieve for genes than for individual transcripts or isoforms, due to the high sequence similarity among groups of isoforms and the non-uniform read coverage resulting from library preparation and sequencing biases [1, 2]. However, gene-level abundance estimation is not without challenges, particularly for groups of genes that share a large fraction of their sequence, which leads to high numbers of multimapping reads [3–5]. Various solutions have been proposed, including grouping together similar genes [4], probabilistic assignment of reads to genes [3] and scoring the genes based on their sequence similarity and number of multi-mapping reads shared with other genes [5].

Despite their higher reliability, gene-level abundances are insufficient for analyses aimed at detecting differences in transcript-level expression or relative isoform usage. Even for studies where the main aim is to detect differential expression on the gene level, incorporating transcript abundances can in some cases improve the inference [2, 6, 7]. As methods for transcript abundance estimation are improving, both in accuracy and speed, it has become increasingly common to estimate abundances of individual isoforms rather than of the gene as a whole, and today a plethora of transcript abundance estimation methods based on various underlying algorithms are available (e.g., [8–17]). Most evaluations of the ability of these methods to accurately estimate transcript abundances have been performed using simulated data, where reads are generated from a known transcriptome [1, 2], or using artificial spike-in sequences [18]. Evaluations have also been done based on the agreement of abundance estimates between replicates [19] or agreement with abundances or abundance ratios derived from other types of data such as exon arrays [20], RT-PCR [21] or 3’ end sequencing [1]. Less is known about the reliability of transcript abundance estimates in real data sets, based on potentially inaccurate or incomplete annotation catalogs, and how to spot unreliably quantified transcripts in a sample-wise manner based on the RNA-seq data alone. A motivating example is illustrated in Figure 1A, showing abundance estimates for the ZADH2 gene in EBV-transformed lymphocytes, as displayed in the GTEx Portal (https://www.gtexportal.org/home/gene/ZADH2, accessed July 19, 2018). This gene has four annotated isoforms, each consisting of two exons and each featuring a unique splice junction (with a shared acceptor site). The top row illustrates the estimated expression of collapsed exons and junctions (with legends to the right), indicating a high expression of the most 5’ exon and the corresponding junction. The alternative exons and junctions have no or very few supporting reads. However, the isoform abundance estimates (lower panel) suggest a different picture, where two of the isoforms whose unique exons and junctions are supported by few reads are assigned the highest expression levels.

**Figure 1:**
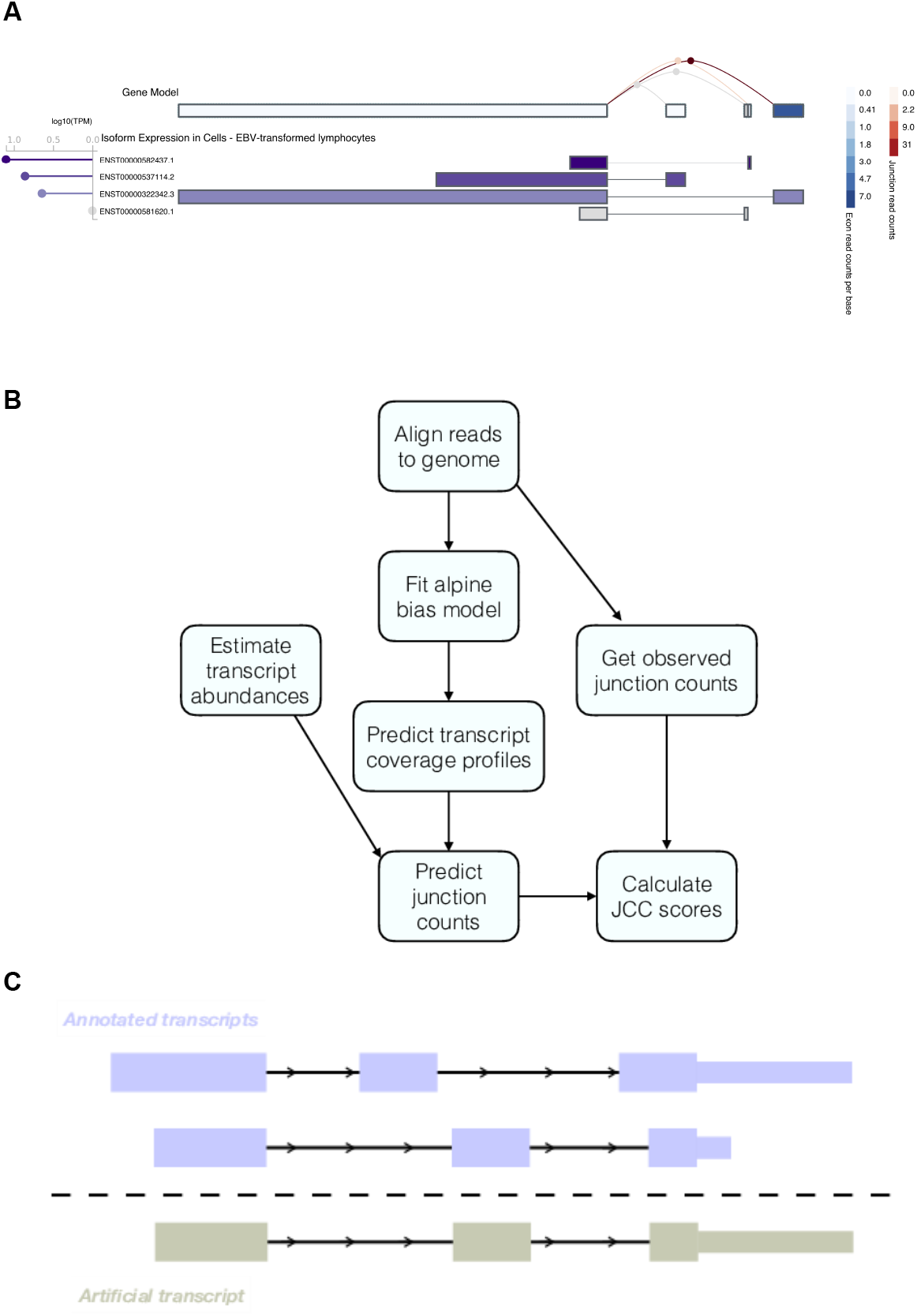
**A**. Example of a gene with inconsistent signals resulting from abundance estimation based on exons, junctions or entire isoforms. The figure was generated in the GTEx Portal (https://www.gtexportal.org/home/gene/ZADH2, accessed July 19, 2018). **B**. Outline of the approach used to calculate the JCC scores. First, reads are aligned to the genome using STAR, and the number of reads observed to span each annotated splice junction is extracted. The aligned reads are also used to fit a fragment bias model using the alpine Bioconductor package, which is then used to predict coverage profiles for all annotated transcripts. The coverage profiles are combined with transcript abundance estimates to obtain the predicted numbers of junction-spanning reads, which are compared to the observed numbers to calculate the JCC score for each gene. **C**. Schematic illustrating the generation of artificial transcripts in the simulated data. In total, artificial transcripts are generated for 4,514 genes, which have multiple annotated 3’UTR of different length (at least 1kb length difference) starting in the same genomic position. For each such gene, two transcripts are selected; one that is annotated with the short 3’UTR and one that is annotated with the long one. The artificial transcript is created by combining the internal structure (all exonic regions except the annotated 3’UTR) of one of the two isoforms with the 3’UTR of the other. In the simulation, all reads from the modified genes are generated from the artificial transcripts.

In this paper, we present the junction coverage compatibility (JCC) score (Figure 1B), which allows automated detection of genes with conflicting indications of isoform abundance. The score can be calculated for any genomic region (e.g., a gene locus), by comparing the observed coverage profile, obtained by aligning the RNA-seq reads to the genome, with the predicted coverage profile derived from estimates of transcript abundances and biases influencing the observed read coverage of a sequenced transcript. In particular, we focus on the number of reads spanning annotated splice junctions in the genomic region of interest. The key assumption behind the JCC score is that with (i) a complete and accurate catalog of reference transcripts, (ii) an accurate estimate of the abundance of each individual transcript, and (iii) knowledge about the biases affecting the prob-ability of a given fragment of a given transcript to be sequenced, the coverage profile prediction obtained by combining these three sources of information for any genomic locus should be close to the observed one. Thus, large deviations between the observed and predicted coverage profiles indicate that the transcript estimates in the region are unreliable, and such regions should be flagged and interpreted with caution in downstream analysis. There can be many reasons behind a region obtaining a high (bad) JCC score, ranging from poor performance of the estimation method, e.g. due to sequence similarity with other parts of the transcriptome, to an incorrect or incomplete annotation catalog, making a correct distribution of the reads between the annotated transcripts in the region impossible.

Using eight transcript abundance estimation methods and two deeply sequenced human RNA-seq data sets, we show that for the majority of the human genes, the junction coverages predicted from the transcript abundances are highly compatible with the observed junction coverages, suggesting overall accurate annotation and transcript abundance estimates. However, a small fraction of the annotated genes show a substantial difference between the predicted and observed junction coverages. For some of these genes, the reason for the incompatibility appears to be an incompletely annotated transcript catalog, and no distribution of the reads among the annotated isoforms would simultaneously give a satisfactory junction coverage compatibility and a good agreement with the annotated UTRs. The uneven read coverage of isoforms also leads to estimation problems, especially for genes with short, poorly covered exons. Using a simulated data set, we show that misannotation of 3’UTRs can lead to unreliable transcript estimates, which is interesting in the light of recent reports showing that the majority of isoform differences between tissues are due to alternative start and end sites and involve untranslated exons [22–24].

## Materials and Methods

### Experimental data and reference annotations

We use two deeply sequenced human polyA+ RNA-seq libraries for our investigations. The first (*Cortex*) contains 117,292,547 paired-end 126nt Illumina reads from a human cerebral cortex sample, and the second (*HAP1*) contains 55,234,720 paired-end 151nt Illumina reads from the HAP1 cell line. Both samples were prepared with the Illumina TruSeq RNA stranded protocol and sequenced at the Functional Genomics Center in Zurich, Switzerland; *Cortex* with a HiSeq 2500 in October 2015 and *HAP1* with a HiSeq 4000 in September 2017. Raw FASTQ files have been uploaded to ArrayExpress (accession number E-MTAB-7089). Most of our analyses are performed using the GRCh38.90 reference annotation from Ensembl [25]. For comparison, we also use the recent CHESS 2.0 reference catalog [26], which was generated by assembling RNA-seq reads from almost 10,000 GTEx samples [27, 28] using StringTie [14]. Based on the original Ensembl gtf file, we generate two additional gtf files, containing flattened exonic regions and intronic regions (regions within a gene locus that are not covered by any exon), respectively, and use featureCounts [29] (from subread [30] v1.6.0) to count the number of reads overlapping these exonic and intronic regions for each gene.

### Simulated data

In addition to the experimental RNA-seq data sets, we generate synthetic data with the aim to better understand the effect of misannotated 3’UTR sequences. From the GRCh38.90 Ensembl annotation, we find 4,514 genes with multiple annotated 3’UTRs starting in the same position, and with length difference exceeding 1kb. For each of these genes, we randomly extract one transcript annotated with the short 3’UTR and one transcript annotated with the long one. We then generate an artificial transcript, consisting of the 5’UTR and coding sequence of one of these two transcripts and the 3’UTR of the other transcript (Figure 1C). For 41 of the 4,514 genes (0.9%), the artificial transcript was identical to an annotated transcript (38 were identical to the transcript providing the 3’UTR, 3 to other isoforms of the gene). These genes were not considered modified. We use the polyester Bioconductor package [31] (v1.16.0) to simulate approximately 1,000 strand-specific read pairs (read length 125nt) from each of the 4,473 remaining artificial transcripts, and a total of 10 million read pairs distributed between 10,000 randomly selected transcripts, not annotated to any of the genes from which the artificial transcripts were generated. The simulated data set is then processed using the original Ensembl GRCh38.90 annotation files (which do not contain the artificial transcripts).

### Transcript abundance estimation

We use eight methods to estimate abundances of the annotated transcripts in each of the two Illumina libraries:

- RSEM. We build an index from the combined cDNA and ncRNA reference fasta files from Ensembl, and estimate transcript abundances with RSEM [9] (v1.3.0), using bowtie [32] (v1.1.2) as the underlying aligner.
- Salmon. We build a transcriptome index from the combined cDNA and ncRNA reference fasta files from Ensembl and run Salmon [16] (v0.11.0) in quasi-mapping mode, incorporating sequence, GC and positional bias correction. We also generate 100 bootstrap samples for estimation of the inferential variance for each transcript. By default, Salmon removes duplicated sequences in the reference catalog, keeping only one representative. In this process, 12,824 transcripts from 4,499 genes were excluded from the Ensembl GRCh38.90 catalog. In a majority of these cases, at least one of the identical sequences can be found on an alternative contig (e.g., in the MHC region). It’s worth noting that these contigs are not included in the primary genome assembly used for the genomic alignments, while the transcripts are contained in the Ensembl transcriptome fasta files. 3,450 of the affected genes did not have any other annotated transcript and were thus completely removed from the annotation catalog.
- SalmonKeepDup. Here we run Salmon with the same settings as above, but retain all duplicated transcript sequences in the catalog (which is an option during Salmon’s indexing step). Since the retained transcripts are sequence-identical, the estimated abundances will be uniformly distributed within groups of identical transcripts.
- kallisto. We build a transcriptome index from the combined cDNA and ncRNA reference fasta files from Ensembl, and run kallisto [15] (v0.44.0) with bias correction activated.
- Hera. The Hera index is built using the reference genome (primary assembly) and the Ensembl gtf file, and Hera (https://github.com/bioturing/hera) (v1.1) is run with default settings.
- HISAT2+StringTie. We build a HISAT2 [33] (v2.1.0) index from the reference genome (primary assembly) and extract the known splice sites using the provided `hisat2_extract_splice_sites.py` script. The reads are aligned to this index with the option `--dta` set and given the known splice sites. Next, we run StringTie [14] (v1.3.3b) without assembly of new transcripts (`-e` option) to get the abundance estimates for the annotated transcripts.
- SalmonSTAR. For this approach we build a transcriptome index from the combined cDNA and ncRNA reference files from Ensembl, and align the reads using STAR [34] (v2.5.3a). We subsequently estimate transcript abundances using Salmon (v0.11.0) in alignment-based mode, incorporating sequence and GC bias correction.
- SalmonCDS. Here, we build the Salmon index using only the explicitly annotated coding sequences from Ensembl, and run Salmon (v0.11.0) in quasi-mapping mode, incorporating sequence, GC and positional bias correction.

### Prediction of expected junction coverage

In order to predict the expected number of reads mapping across each junction, given estimates of the transcript abundances, we first fit a fragment-level bias model using the `alpine` Bioconductor package [35] (v1.2.0). The bias model is fit for each library separately, using a set of single-isoform genes with length between 600 and 7,000 bp and between 500 and 10,000 assigned reads. The `alpine` bias model included random hexamer bias, fragment GC bias, positional bias along the transcript, and the fragment length distribution. After fitting the bias model, we use it to obtain a predicted coverage of each base pair in each annotated transcript using the fitted parameters for these four factors. For transcripts where the prediction fails (e.g., transcripts shorter than the estimated fragment length and transcripts with no overlapping reads), we assume a uniform coverage rather than excluding them from subsequent analysis steps. Next, we rescale the predicted base pair coverages by dividing with their total sum and multiplying with the average fragment length and the estimated transcript counts from each of the transcript abundance estimation methods, in order to get an estimate of the number of reads predicted to cover each position on the transcript. We also extract the position of annotated splice junctions within each transcript, and the predicted coverage at the base just before an annotated junction is used as the predicted number of reads from that transcript that align across the junction. Finally, we sum the predicted number of junction-spanning reads for each junction across all transcripts, in a strand-aware fashion (since the libraries are stranded) in order to get the total number of reads predicted to span any given junction.

### Observed junction coverage

The observed junction coverage (the number of reads mapping across a given junction) is obtained using STAR [34] (v2.5.3a). We build an index using the reference genome (primary assembly) and the Ensembl gtf file, and align the reads with default settings. The number of uniquely mapping and multimapping reads spanning each annotated junction are extracted from the SJ.out.tab output file from the STAR alignment.

### The junction coverage compatibility score

To quantify the level of agreement between the predicted junction coverages based on any of the transcript abundance estimation methods and the observed number of junction reads from STAR, we define a family of gene-wise junction coverage compatibility (JCC) scores, parametrized by two arguments: a weighting function *g* and a scaling indicator *β* (see below). For a given *g* and *β*, the JCC score for gene *i* is defined by

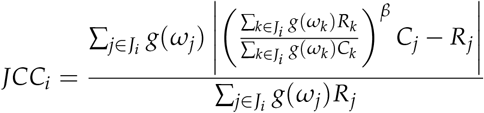

where *J_i_* denotes the set of junctions annotated to gene *i* (some junctions are annotated to transcripts from multiple genes, in which case they are included for all of them), *R_j_* is the observed number of uniquely mapping reads spanning junction *j* (obtained from STAR) and *C_j_* is the predicted number of reads spanning junction *j* based on the bias model from alpine and the transcript abundances from a given method. Multimapping reads (from STAR) cause problems in the score calculation since it is not clear how to assign them to junctions, and thus the contribution of a junction is weighted by *g*(*ω_j_*), where *g*: [0, 1] ↦ [0, ∞) is a non-negative function and *ωj* is the fraction of reads spanning junction *j* that are uniquely mapping.

Overall differences in the number of reads assigned to gene *i* by transcript abundance estimation compared to junction counts can induce large differences between *C_j_* and *R_j_* even if their relative coverage patterns are similar. The same is true if there is a large fraction of multimapping reads, which are being accounted for in the predicted transcript abundances but not in the observed junction coverages. To account for this, we include an optional scaling of the predicted coverages to have the same (weighted) sum as the observed coverages. This is represented by the *β* parameter - if this is 0, no scaling is done, and if it is 1, the values are scaled. In this study, we set *β* = 1, and let

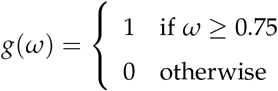

i.e., a step function that implies that we only allow junctions with more than 75% uniquely aligning reads to contribute to the JCC score calculations.

With *β* = 1, the JCC score for a gene takes values between 0 and 2. A low score means that the predicted junction coverages, given the abundance estimates for the transcripts in gene *i*, are compatible with the observed number of reads mapping across the junctions, while a high score indicates that for at least one junction, the predicted number of junction-spanning reads does not match with the observed number.

### Code availability

All code used to perform the analyses is available from https://github.com/csoneson/annotation_problem_txabundance.

## Results

### Predicted transcript coverage patterns agree well between samples

The prediction of the transcript coverage profiles by alpine is a crucial step in the calculation of the JCC score. It is done separately for each of the two Illumina libraries, in order to account for any sample-specific biases. Of the 200,310 annotated transcripts in the Ensembl gtf file, the prediction of the coverage pattern by alpine returned an expected error for 29,342 (14.6%) in the *HAP1* sample and 13,906 (6.9%) in the *Cortex* sample, almost exclusively due to transcripts being shorter than the respective fragment lengths. The prediction returned NULL for 23,028 (11.5%) transcripts in the *HAP1* sample and 11,941 (6.0%) in the *Cortex* sample that did not have any overlapping reads. For these transcripts we impose a uniform coverage, rather than excluding them from subsequent calculations.

Overall, we observe a high correlation between the predicted coverage profiles in the two libraries (Supplementary Figure 1), indicating that they share many of the biases, despite coming from different cell types and being prepared and sequenced almost two years apart on different sequencing machines. The coverage prediction is the single most time-consuming step of the JCC score calculation, and the high correlation even between such different libraries suggests that in a specific study, the prediction may not need to be done separately for each individual sample, which can reduce the run time considerably. Run time can also be reduced by limiting the coverage prediction and subsequent analysis to transcripts from a subset of the genes that are of particular interest in a given situation.

### Most predicted junction coverages are consistent with the observed coverages

Using the approach described in the Methods section, we obtain the number of uniquely mapping reads observed to span each annotated junction as well as the number predicted to span each junction given each set of transcript abundance estimates. Comparing the predicted junction coverages *C_j_* to the observed ones *R_j_* across all annotated junctions shows a generally high correlation for all abundance estimation methods (Figure 2A, left column), suggesting that in most genomic loci, the annotated transcript structure is compatible with the observed read alignments, and that the approach we use to predict junction coverages based on transcript abundances is valid. Scaling the predicted junction coverages within each gene, corresponding to setting *β* = 1 in the subsequent JCC calculation and thereby focusing more on the relative junction coverages within a gene rather than the overall abundance of the gene, increases the correlation for all methods (Figure 2A, right column). The largest discrepancies between observed and predicted junction coverages are seen for SalmonCDS, indicating that on a global scale, only considering annotated coding sequences discards relevant information about transcript abundances. We also note that there is a set of junctions with a low fraction of uniquely mapping reads (Figure 2A, marked in red) for which the predicted number of spanning reads is considerably higher than the observed number of uniquely mapping junction reads. Since these discrepancies do not represent a failure of the annotation system or transcript abundance estimation method, but rather an inability to place reads in a unique genomic position, we downweight the influence of these junctions on the JCC score via the *g*(*ω*) function, as described in the Methods section.

**Figure 2:**
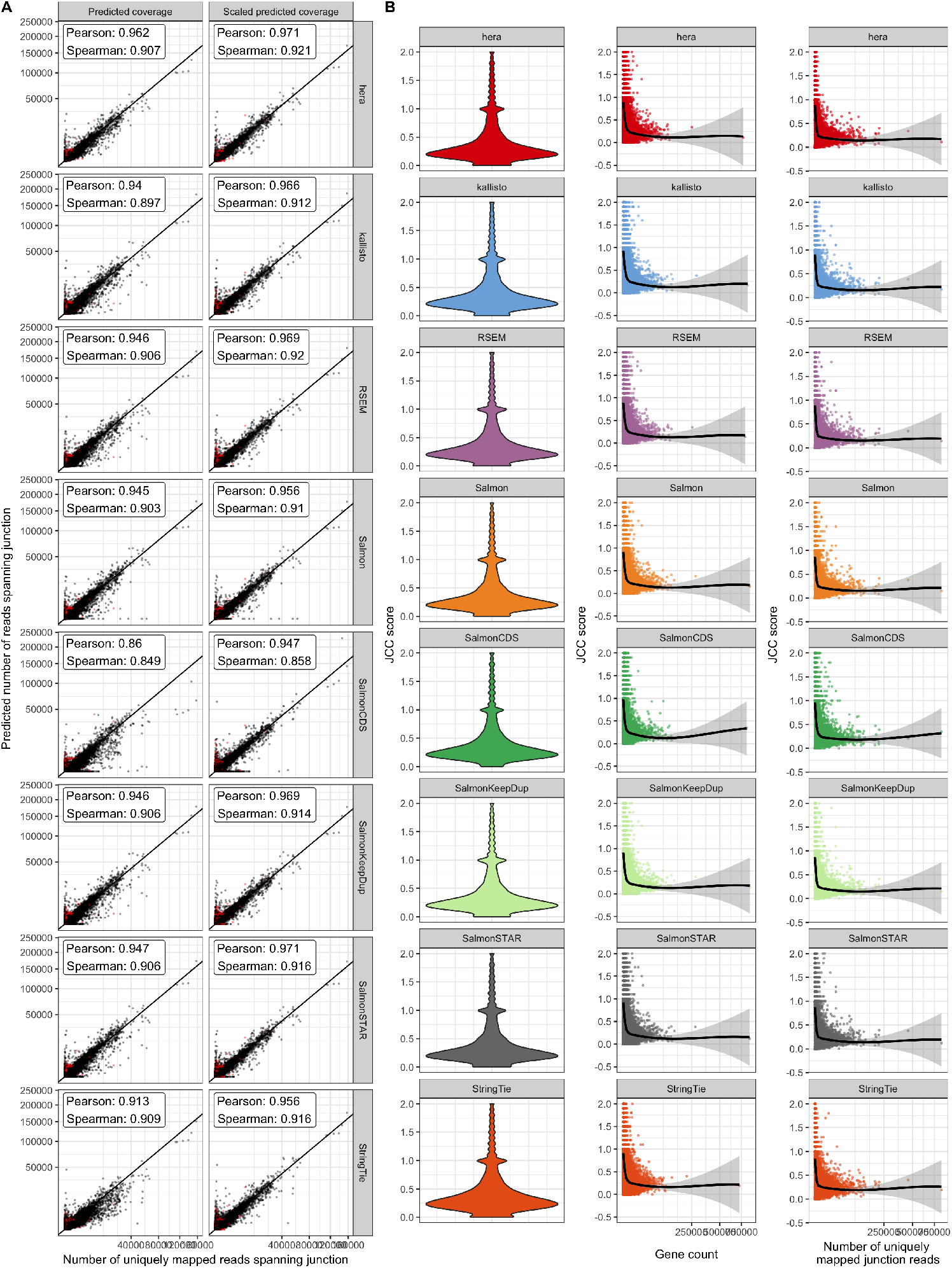
**A**. Correlation between observed and predicted number of reads spanning each junction for the *HAP1* sample. The left column (“Predicted coverage”) shows the actual number of reads predicted by `alpine`, while the predicted values in the right column (“Scaled predicted coverage”) are scaled to sum to the same number as the observed number of uniquely mapping junction reads within each gene. Scaling improves the correlation between observed and predicted junction coverages for all included methods. Axes are square-root transformed for better visualization. Red points indicate junctions where less than 75% of the spanning reads are uniquely mapping. **B**. Overall distribution of the gene-wise JCC scores for each method in the *HAP1* sample, as well as the association between the JCC score and the total number of reads for the gene and the number of uniquely mapped junction reads in the gene.

### Most genes show high compatibility between observed and predicted junction coverages

After investigating the concordance between observed and predicted coverages for individual junctions, we next calculate the JCC score for each annotated gene. With the exception of SalmonCDS (which is using a reference annotation in which many transcripts and genes are missing since they don’t have an explicitly annotated coding sequence), we are able to calculate a valid JCC score for around 16,500 genes in the *HAP1* library, and just over 20,000 genes in the *Cortex* library (Supplementary Figure 2). Among the genes for which the score can not be calculated, most are not expressed (predicted total abundance of all isoforms equal to 0), while a smaller fraction either are expressed but lack junctions, or contain junctions but have no or too few uniquely mapping junction-spanning reads to calculate the score.

Investigating the overall distribution of valid JCC scores shows that for most genes, the score is low (below 0.5), confirming the previous observation that for most of the genes, the junction coverage pattern induced by the estimated transcript abundances agrees well with the observed junction coverages (Figure 2B, left column). Similar distributions are seen for all included methods. Most of the very high scores are obtained for genes with low abundance and few uniquely mapped reads spanning any of the junctions (Figure 2B). The high score for these genes may be driven largely by shot noise, and may improve with even higher sequencing depth. Moreover, lowly expressed genes are typically excluded in practical analyses of RNA-seq data such as differential expression analyses. Thus, to illustrate the behaviour of the JCC score, in the following analyses we focus on genes with at least 25 reads mapping uniquely across any of its junctions.

### Examples of genes with high JCC score

In order to exemplify the types of deviating patterns resulting in high JCC scores, we consider some of the genes that are assigned high scores (JCC ≥ 0.6) with all the transcript abundance methods (except SalmonCDS, since it is based on a different set of reference transcripts and does not represent a typical or recommended way of performing transcript abundance estimation). Furthermore, we limit the investigation to genes with at least 25 uniquely mapped junction-spanning reads, at least 75% of the junction-spanning reads mapping uniquely and an intron/exon read count ratio below 0.1. These strict filtering criteria are satisfied by 146 genes in the *Cortex* library and 56 genes in the *HAP1* library. 17 of the genes passed the filters in both libraries. One of these genes is ZADH2 (Figure 3, other examples in Supplementary Figures 3-4). ZADH2 has four annotated transcripts, each consisting of two exons and one junction, and no junction is shared between transcripts. Most transcript abundance estimation methods distribute the estimated abundance between two or three of these isoforms. However, only one of the four annotated junctions has any observed spanning reads, which suggests that only the corresponding transcript (ENST00000322342) is indeed present. This leads to a large discrepancy between the observed and predicted junction coverages (for all abundance estimation methods), and hence a large JCC score. For this gene, a possible explanation for the discrepancy is that the coverage of the 5’ end of the transcripts is weak, but for a reason not captured by the alpine bias model, implying that the 3’ end, which is longer and shows a higher coverage, will dominate the abundance estimation. Uneven coverage in this region can therefore bias the abundance estimation towards one or the other transcript. As illustrated in Figure 1A, a similar behaviour can be seen also in the GTEx data (accessed via the GTEx Portal).

**Figure 3:**
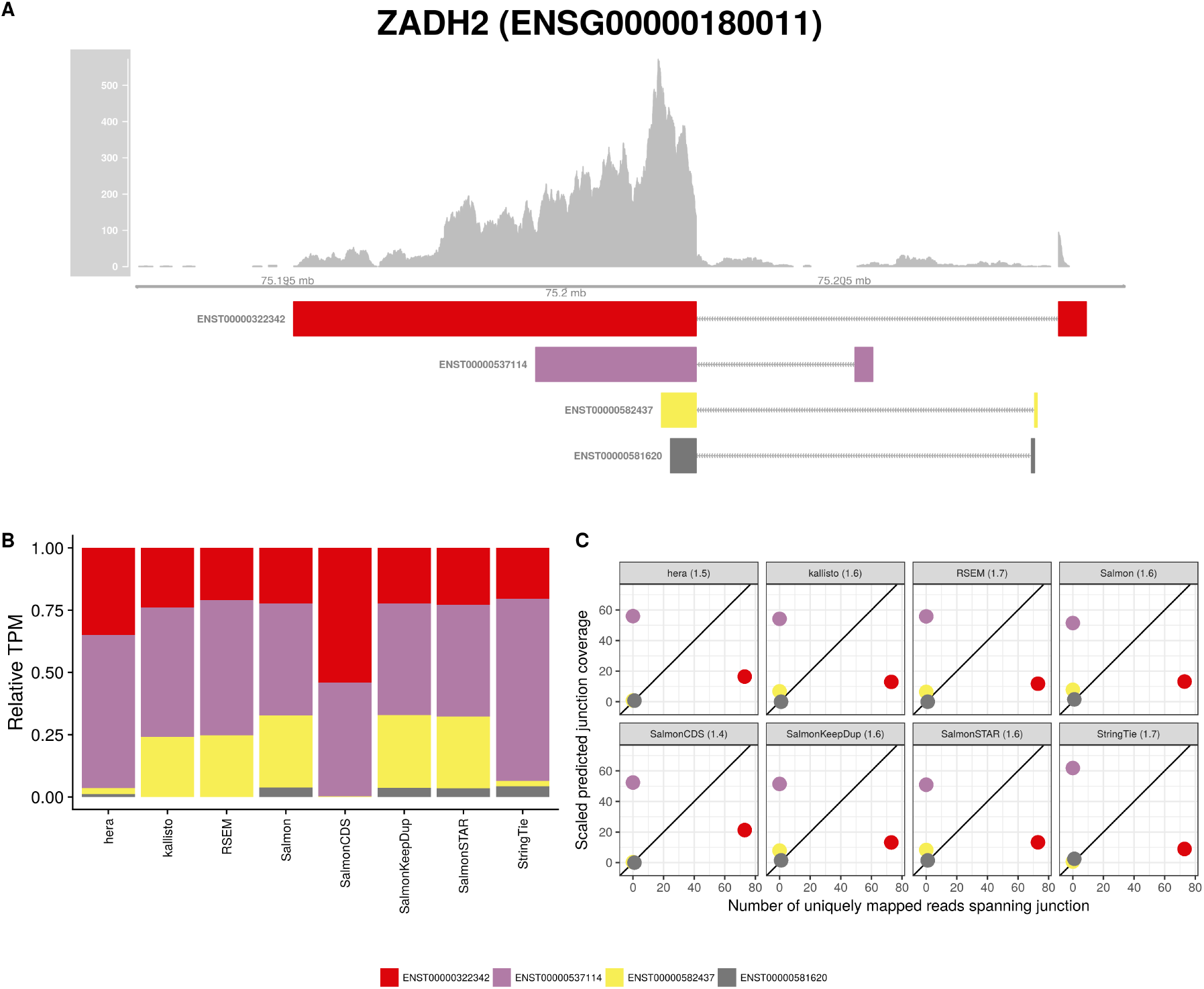
**A**. Observed coverage profile and annotated gene model for the ZADH2 gene in the *HAP1* library. Different annotated transcripts are shown in different color. **B**. Relative TPM estimates for the annotated transcripts from each of the eight transcript abundance estimation methods. **C**. Observed number of uniquely mapping junction-spanning reads (*x*) and scaled predicted junction coverages (*y*) based on transcript abundance estimates from each of the eight methods. Each circle corresponds to an annotated junction and is colored according to the set of transcripts that it is annotated to. The JCC scores for this gene based on the respectively abundances are indicated in the panel headers.

Similar observations can be made for many of the other selected example genes (Supplementary Figures 3-4). For many of the high-scoring genes, no distribution of the reads between the annotated transcripts would lead to compatible coverage of both the internal structure (junctions) and the UTRs. For other genes with high scores, we observe a read coverage that is largely incompatible with the annotated exons (e.g., Supplementary Figure 5). In particular, these genes have large numbers of intronic reads, which can not be accounted for by annotated transcripts. This suggests that high scores can often be attributed to incorrectness or incompleteness of the annotation catalog with respect to the observed reads, rather than poor performance of the abundance estimation methods. Regardless of the cause, however, the resulting abundances are unreliable and should be interpreted with caution in downstream analyses.

### JCC scores are not strongly associated with inferential variability

Several isoform abundance estimation methods allow assessment of the variability of the resulting expression levels via some form of (re)sampling [9, 10, 15, 16, 36, 37]. In order to compare the uncertainties picked up by the JCC score to those represented in these inferential variances, we perform 100 bootstrap runs using Salmon, and estimate the coefficient of variation of the bootstrapped counts both on the transcript level and after aggregating the transcript counts on the gene level. For the evaluation, we consider only genes with at least 25 uniquely mapping junction-spanning reads, and each individual transcript is assigned the JCC score of the corresponding gene. Overall, the association between the inferential coefficient of variation and the JCC score is weak in both libraries, on both the transcript and gene level (Supplementary Figure 6). Thus, the two scores measure different types of uncertainties; while the bootstrap variability may capture assignment uncertainty caused by shared sequence features among transcripts, it will not in general pick up inconsistencies due to misannotation, which are targeted by the JCC score.

### JCC scores are overall similar between methods

Since the JCC score is obtained by combining a set of estimated transcript coverage profiles with transcript abundance estimates, using different transcript abundance methods for the latter leads to different sets of scores. We calculate JCC scores using transcript abundance estimates from eight different methods, and the results shown above illustrate that all of them suffer from problems induced by misannotated or missing transcripts, leading to predicted junction coverages that are incompatible with the observed ones. To further investigate the similarities between the methods, we calculate correlation coefficients between the scores obtained by each method pair, using only genes with at least 25 uniquely mapping junction-spanning reads (Supplementary Figures 7-9). As expected, the correlation is overall very high, and the most deviating scores are obtained with SalmonCDS, which uses a different set of reference sequences than the other methods, and StringTie. On average, both SalmonCDS and StringTie give higher scores than the remaining methods (Supplementary Figure 9B).

### The choice of reference annotation affects the JCC score distribution

All analyses so far have been performed using the Ensembl GRCh38.90 annotation. To investigate the impact of the choice of reference annotation on the JCC scores, we estimate bias models and predict transcript coverage profiles also for all transcripts in the CHESS 2.0 catalog [26]. We estimate corresponding transcript abundances with Salmon and kallisto, and count junction-spanning reads for each annotated junction with STAR. The CHESS catalog is obtained by assembling reads from almost 10,000 GTEx samples, and contains a larger number of transcripts (annotated to a smaller number of genes) than the Ensembl catalog (Supplementary Table 1). The CHESS genes are all annotated with a unique CHESS identifier, but a mapping to Entrez IDs is provided wherever possible. For comparison with our other results, we convert the Entrez IDs to Ensembl IDs using the `org.Hs.eg.db` Bioconductor package v3.6.0 (in this way, unique Ensembl IDs are obtained for for 22,262/42,881=51.9% of the genes). Considering only genes that are shared between the two annotation catalogs, it is clear that there is a substantial difference between the scores assigned to an individual gene using the two annotations (Figure 4A), although the overall distribution of scores is largely similar (Figure 4B). Neither annotation catalog is consistently leading to lower scores than the other (Figure 4C), but there are genes with substantially lower scores with each of the two annotations compared to the other.

**Figure 4:**
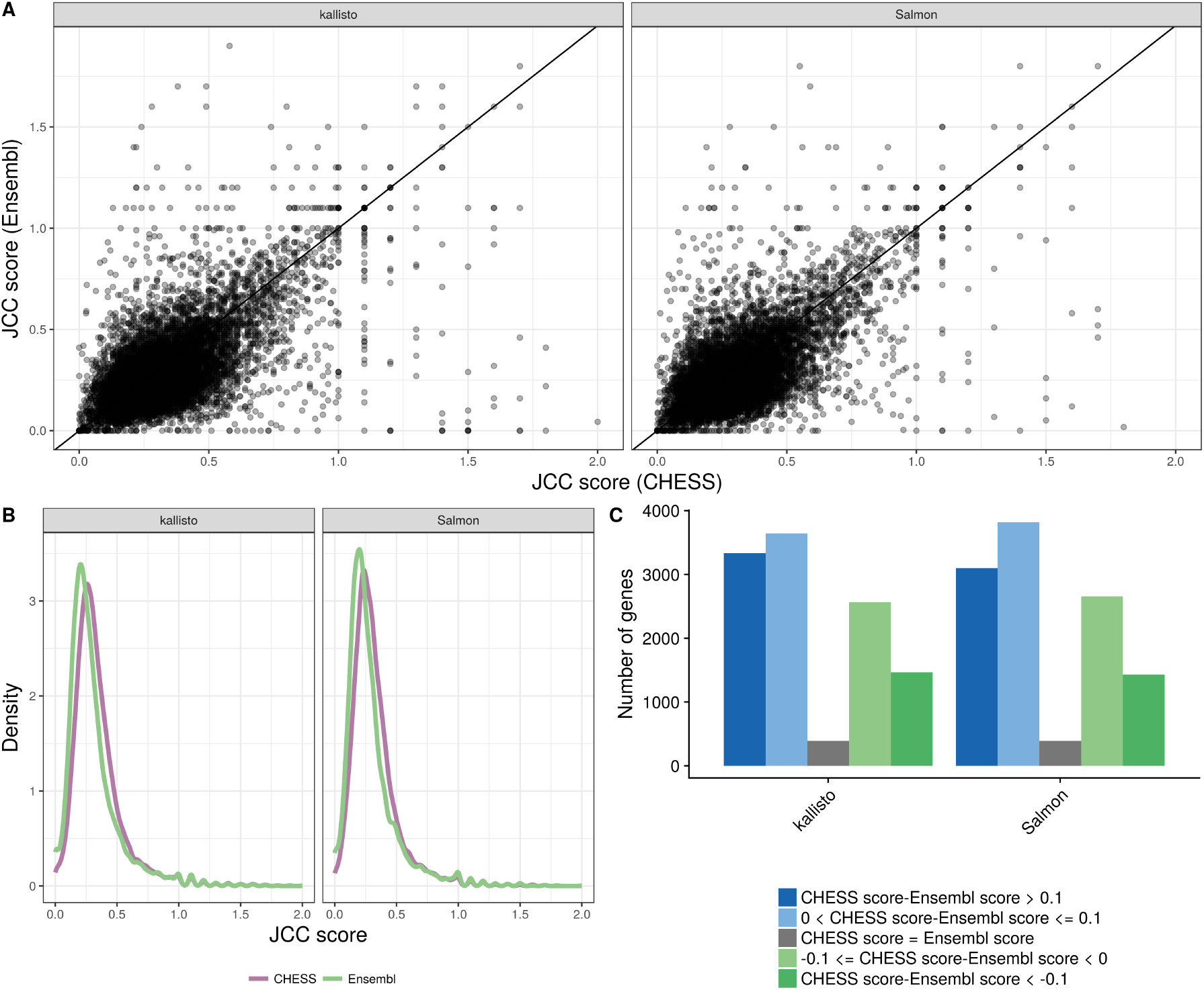
Comparison between scores obtained with the Ensembl GRCh38.90 annotation and the CHESS 2.0 annotation, for the *HAP1* sample. **A**. Correlation between scores obtained with the CHESS annotation (x) and the Ensembl annotation (y), for all the shared genes (genes with an assigned Ensembl ID in the CHESS catalog), with at least 25 uniquely mapping junction-spanning reads and at least 75% of the junction-spanning reads mapping uniquely with both annotations. **B**. Distribution of JCC scores for all genes with at least 25 uniquely mapping junction-spanning reads and at least 75% of the junction-spanning reads mapping uniquely, in the respective annotation catalogs. **C**. The number of genes shared between the two annotation catalogs for which the CHESS annotation results in a higher, lower or equal score compared the Ensembl annotation. Blue bars represent genes for which scores based on the CHESS annotation are higher (worse) than those based on the Ensembl annotation, and green bars represent the opposite situation.

In addition, we investigate the effect of quantifying transcript abundances using a dataset-specific catalog of transcripts, obtained by running StringTie (without the -e argument) on each of the two Illumina libraries. The resulting gtf file contains many new transcripts, and many annotated transcripts from the Ensembl catalog are removed (Supplementary Table 1). We apply a subset of the abundance estimation methods to the respective StringTie annotations, and compare JCC scores across all genes present in both the StringTie and Ensembl catalogs. Also in this case, no annotation consistently lead to lower scores than the other, but there is a larger fraction of genes that show lower scores with the sample-specific StringTie-assembled annotation (Supplementary Figure 10).

### Misannotated 3’UTRs strongly affect the abundance estimates

To investigate the effect of misannotated or missing 3’UTRs on the transcript abundance estimates, and consequently the JCC score, in more detail, we used synthetic data. For each of 4,514 annotated genes, we generated an artificial transcript consisting of the coding sequence of one isoform and the 3’UTR of another isoform from the same gene. The two contributing isoforms were selected in such a way that one was annotated with a short 3’UTR, and the other with a long 3’UTR (with a length difference of at least 1kb) starting in the same genomic location. As expected, for genes where the isoform with the long 3’UTR was selected to contribute the 3’UTR to the artificial transcript, a large fraction of the final artificial transcript consists of the 3’UTR, while the fraction is much smaller if the 3’UTR was chosen from the isoform with the short 3’UTR (Supplementary Figure 11).

For the modified genes, reads are simulated only from the artificial transcript. We also simulate reads from a random selection of unmodified transcripts. As expected, the JCC scores for the genes with modified transcripts are generally higher than those for the genes without any modified transcripts, where the reads are simulated from the correct annotation catalog. The distribution of scores for the latter set of genes can be seen as a “baseline distribution” of scores that we can expect for reasons unrelated to annotation and sequencing artifacts (e.g., sequence similarity causing problems for abundance estimation methods). Furthermore, the JCC score is generally higher for genes where a larger fraction of the artificial transcript is made up of the 3’UTR (Supplementary Figure 12). Focusing only on the genes with modified transcripts, we calculate the similarity between the artificial transcript and all annotated transcripts from the same gene. The similarity is defined by the Jaccard index of the nucleotide positions covered by the two compared transcripts. We stratify the genes based on whether the most similar transcript to the artificial transcript is the one that contributed the internal structure, the one that contributed the 3’UTR, or another one of the annotated transcripts. For most abundance estimation methods, the annotated transcript that is most similar to the artificial transcript (from which the reads were generated) is also assigned the highest expression estimate (Figure 5). The exceptions are SalmonCDS and StringTie, which both generally assign the highest abundance to the transcript that is most similar to the artificial transcript in terms of the internal structure, rather than based on overall similarity.

**Figure 5:**
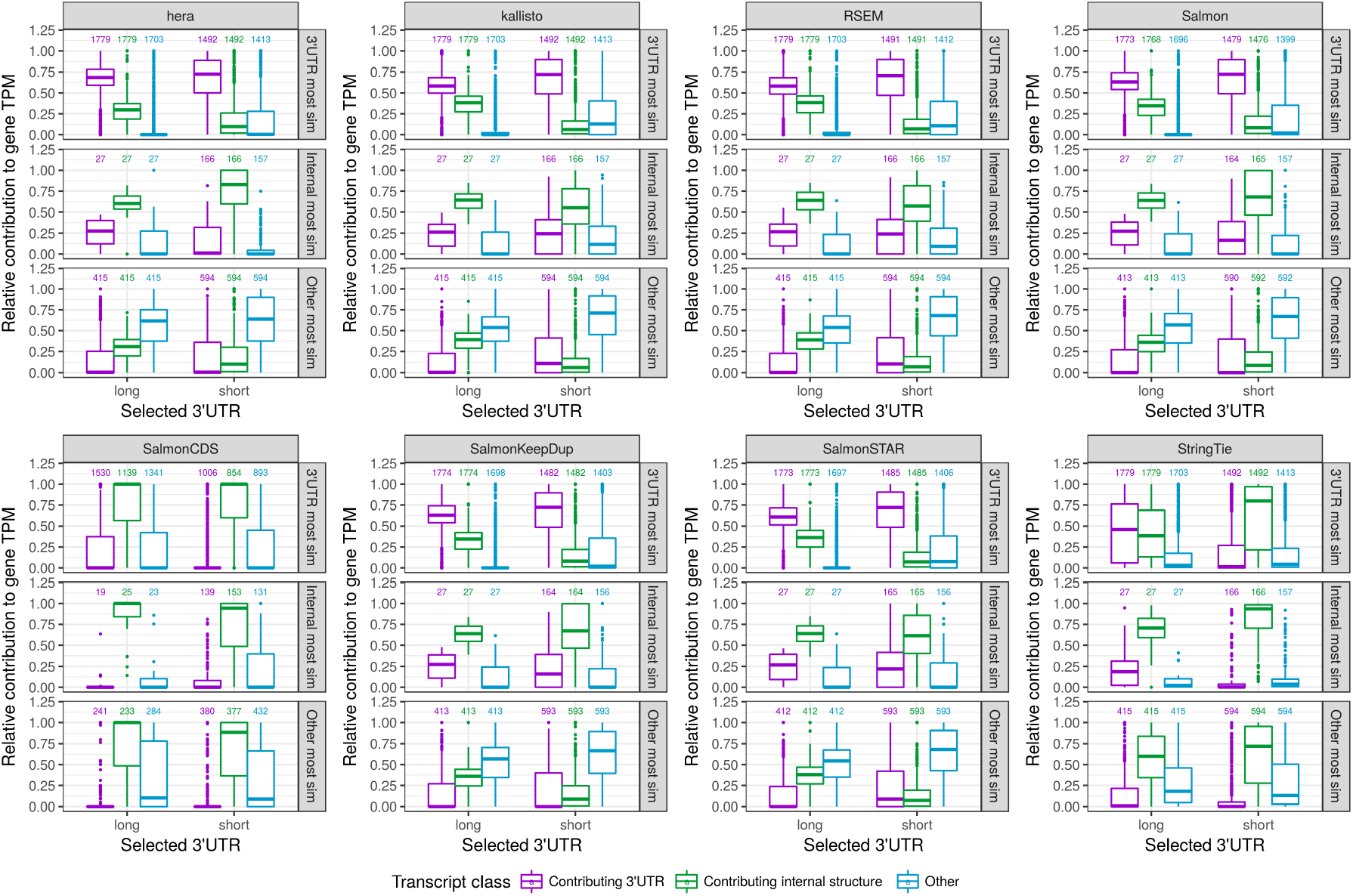
Relative transcript abundances for modified genes, with each of the eight transcript abundance estimation methods. Genes are stratified (vertically) based on whether the transcript that is most similar (by Jaccard index of covered nucleotides) to the artificial transcript is the one contributing the 3’UTR, the one contributing the internal structure, or another isoform of the gene (see Figure 1C). For each gene we calculate the relative abundance of the transcript contributing the 3’UTR, the one contributing the internal structure, and all other isoforms of the gene combined (indicated with color). Finally, genes are stratified (horizontally) based on whether the artificial transcript contains the long or short variant of the 3’UTR. Generally, most methods assign the highest abundance to the transcript that is most similar to the artificial transcript from which the reads were generated, with the exception of SalmonCDS and StringTie, which assign higher abundances to the transcripts that are most similar to the artificial transcript in terms of the internal structure. The numbers above the boxplots indicate the number of genes in each category.

## Discussion and Conclusions

We have described the junction coverage compatibility (JCC) score and shown how it can be used to identify genes or genomic regions where junction coverage patterns predicted from estimated transcript abundances are incompatible with those observed after alignment of the RNA-seq reads directly to the genome. By using the RNA-seq data to obtain two estimates of the number of reads mapping across each splice junction, we can create an internal validation system, thereby circumventing the need for an external data set or additional replicates for evaluation of transcript abundance estimation accuracy. A high score, indicating poor compatibility between the junction coverages estimated from the transcript abundance estimates and the observed junction coverages, can be caused, e.g., by inaccurate transcript abundance estimates (e.g., for transcripts that share large parts of their sequence with other transcripts), or by an incomplete or incorrect transcriptome annotation. Regardless of the underlying cause, such genes should be flagged in downstream analyses and the estimated transcript abundances interpreted with caution. We note that the results were overall similar for all the eight transcript abundance estimation tools used in the study, representing alignment-free methods as well as methods relying on either genome or transcriptome alignments.

The chosen reference annotation can have a large effect on the resulting JCC scores, as seen here by comparing the scores obtained using the Ensembl annotation to those based on the CHESS 2.0 annotation. In addition, using StringTie to assemble missing transcripts led to improved scores for a large number of genes, and a worse score for a smaller number of genes. As recommended^1^, we used the primary genome assembly from Ensembl for aligning the reads to the genome. However, the transcriptome fasta files from Ensembl, which were used as the basis for abundance estimation by Salmon, SalmonKeepDup, kallisto, RSEM and SalmonSTAR, contain transcripts from alternative contigs that are not included in the primary genome assembly. Many of these transcripts are identical or very similar to transcripts annotated to the primary chromosomes. While this represents the typical use of these alignment files for alignment and transcript abundance estimation, it may lead to problems for the correct assignment of the reads to transcripts, and as a consequence, for the calculation of the JCC scores. Keeping only one representative of duplicate transcript sequences (the default behaviour of Salmon) can lead to both better abundance estimates and improved agreement between predicted and observed junction coverages, under the assumption that the correct transcript location is retained. Of course, determining the true location of origin of a given transcript can be highly non-trivial, but would be an interesting direction for future research.

One limitation of the presented family of JCC scores is that they can not be calculated for genes that do not have annotated junctions, or that do not have reads spanning junctions. A solution to this could be to compare the predicted and observed coverage profiles of the entire genomic locus rather than just the junctions. However, multimapping reads will still pose a problem for the comparison, and positions with a large fraction of multimapping overlapping reads should be downweighted in the score. In general, the approach we propose is not limited to junction coverages, and could be extended to, e.g., disjoint exon bins. The requirement is that we can observe the coverage pattern of the features of interest from the genome alignment, and predict it from the alpine bias models and the estimated transcript abundances. In addition, while we use the weighting function *g*(*ω*) to downweight the influence of junctions with a large fraction of multi-mapping reads, it can be used more generally to assign weights to junctions based on any characteristics affecting our confidence in the observed read coverages.

Our evaluations are based on the assumption that we are interested in obtaining and using transcript abundance estimates. Other quantification approaches, for example, those focusing on disjoint exon bins [38] or transcript equivalence classes [39] have been suggested, and the resulting counts may in themselves be less sensitive to uncertainties in the reference transcript catalog. However, a post-processing step is required in order to interpret the results in terms of known transcripts, and during this step, misannotated transcripts can still lead to erroneous conclusions.

Using simulated data, we observed that compared to the other abundance estimation methods, StringTie appeared to focus more on matching the internal structure than the 3’UTR when assigning abundances to transcripts. This implies that in situations where the 3’UTR annotation is unclear, StringTie can help assigning the reads to the transcript that is most similar with respect to the more unambiguous part of the transcript structure. However, it could potentially also make it more difficult to identify differences in transcript composition between tissues, since these have been shown to be predominantly differences in transcription start and end sites [24].

Our results show that for the vast majority of the human genes, the junction coverage patterns implied by the estimated transcript abundances in our data sets agree well with the observed ones, indicating that the reference annotation as well as the transcript abundance estimates for these genes are likely to be reliable. However, for each transcript abundance estimation method a small number of genes obtained a high JCC score, suggesting unreliably quantified isoforms. These genes should be treated with care in any downstream analyses, or be investigated further for an improved transcriptome annotation.

## Acknowledgements

The authors would like to thank the members of the Robinson, von Mering and Baudis groups at the University of Zurich for helpful discussions.

## Footnotes

1 e.g., https://github.eom/alexdobin/STAR/blob/2.5.3a/doc/STARmanual.pdf

